# Neutralizing antibody and soluble ACE2 inhibition of a replication-competent VSV-SARS-CoV-2 and a clinical isolate of SARS-CoV-2

**DOI:** 10.1101/2020.05.18.102038

**Authors:** James Brett Case, Paul W. Rothlauf, Rita E. Chen, Zhuoming Liu, Haiyan Zhao, Arthur S. Kim, Louis-Marie Bloyet, Qiru Zeng, Stephen Tahan, Lindsay Droit, Ma. Xenia G. Ilagan, Michael A. Tartell, Gaya Amarasinghe, Jeffrey P. Henderson, Shane Miersch, Mart Ustav, Sachdev Sidhu, Herbert W. Virgin, David Wang, Siyuan Ding, Davide Corti, Elitza S. Theel, Daved H. Fremont, Michael S. Diamond, Sean P.J. Whelan

## Abstract

Antibody-based interventions against SARS-CoV-2 could limit morbidity, mortality, and possibly disrupt epidemic transmission. An anticipated correlate of such countermeasures is the level of neutralizing antibodies against the SARS-CoV-2 spike protein, yet there is no consensus as to which assay should be used for such measurements. Using an infectious molecular clone of vesicular stomatitis virus (VSV) that expresses eGFP as a marker of infection, we replaced the glycoprotein gene (G) with the spike protein of SARS-CoV-2 (VSV-eGFP-SARS-CoV-2) and developed a high-throughput imaging-based neutralization assay at biosafety level 2. We also developed a focus reduction neutralization test with a clinical isolate of SARS-CoV-2 at biosafety level 3. We compared the neutralizing activities of monoclonal and polyclonal antibody preparations, as well as ACE2-Fc soluble decoy protein in both assays and find an exceptionally high degree of concordance. The two assays will help define correlates of protection for antibody-based countermeasures including therapeutic antibodies, immune γ-globulin or plasma preparations, and vaccines against SARS-CoV-2. Replication-competent VSV-eGFP-SARS-CoV-2 provides a rapid assay for testing inhibitors of SARS-CoV-2 mediated entry that can be performed in 7.5 hours under reduced biosafety containment.

## INTRODUCTION

Severe acute respiratory syndrome coronavirus 2 (SARS-CoV-2) is a positive-sense, single-stranded, enveloped RNA virus that was first isolated in Wuhan, China in December, 2019 from a cluster of acute respiratory illness cases (Guan et al., 2020). SARS-CoV-2 is the etiologic agent of coronavirus disease 2019 (COVID-19), which as of May 16, 2020 has more than 4.5 million confirmed cases causing 309,000 deaths. Virtually all countries and territories have been affected, with major epidemics in Central China, Italy, Spain, France, Iran, Russia, the United Kingdom, and the United States. SARS-CoV-2 is thought to be of zoonotic origin and is closely related to the original SARS-CoV (Zhang et al., 2020; Zhou et al., 2020). Most cases are spread by direct human-to-human transmission, with community transmission occurring from both symptomatic and asymptomatic individuals (Bai et al., 2020). This has resulted in a global pandemic with severe economic, political, and social consequences. The development, characterization, and deployment of an effective vaccine or antibody prophylaxis or treatment against SARS-CoV-2 could prevent morbidity and mortality and curtail its epidemic spread.

The viral spike protein (S) mediates all steps of coronavirus entry into target cells including receptor binding and membrane fusion (Tortorici and Veesler, 2019). During viral biogenesis the S protein undergoes furin-dependent proteolytic processing as it transits through the trans-Golgi network and is cleaved into S1 and S2 subunits that function in receptor binding and membrane fusion, respectively (Walls et al., 2020). Angiotensin-converting enzyme 2 (ACE2) serves as a cell surface receptor (Letko et al., 2020; Wrapp et al., 2020) for SARS-CoV-2, and productive infection is facilitated by additional processing of S2 by the host cell serine protease TMPRSS2 (Hoffmann et al., 2020).

Laboratory studies of SARS-CoV-2 require biosafety level 3 (BSL3) containment with positive-pressure respirators. Single-round pseudotyped viruses complemented by expression of the SARS-CoV-2 S protein *in trans* serve as biosafety level 2 (BSL2) surrogates that can facilitate studies of viral entry, and the inhibition of infection by neutralizing antibodies and other inhibitors (Hoffmann et al., 2020; Lei et al., 2020; Ou et al., 2020). Such pseudotyping approaches are used routinely by many laboratories for other highly pathogenic coronaviruses including SARS-CoV and MERS-CoV (Fukushi et al., 2006; Fukushi et al., 2005; Giroglou et al., 2004; Kobinger et al., 2007). Viral pseudotyping assays are limited by the need to express the glycoprotein *in trans* and preclude forward genetic studies of the viral envelope protein. Expression of the glycoprotein is often accomplished by plasmid transfection, which requires optimization to minimize batch variation. Assays performed with such pseudotyped viruses rely on relative levels of infectivity as measured by a reporter assay without correlation to an infectious titer. It also is unknown as to how the display of S proteins on a heterologous virus impacts viral entry, antibody recognition, and antibody neutralization compared to infectious coronavirus. This question is important because neutralization assays are used to establish correlates of protection for vaccine and antibody-based countermeasures, and most manufacturers lack access to high-containment laboratories to test antibody responses against highly pathogenic coronaviruses including SARS-CoV-2.

Here, we developed a simple and robust BSL2 assay for evaluating SARS-CoV-2 entry and its inhibition by antibodies. We engineered an infectious molecular clone of vesicular stomatitis virus (VSV) to encode the SARS-CoV-2 S protein in place of the native envelope glycoprotein (G) and rescued an autonomously replication-competent virus bearing the spike. Through passage of VSV-eGFP-SARS-CoV-2, we selected a gain-of-function mutation in S that allowed more efficient viral propagation yielding titers of > 1 x 10^8^ plaque-forming units (PFU)/ml. We characterized this variant with respect to inhibition by soluble human ACE2-Fc and monoclonal and polyclonal antibodies from humans and compared those results to neutralization tests with a clinical isolate of SARS-CoV-2. These studies demonstrate that a recombinant VSV expressing SARS-CoV-2 S behaves analogously to a clinical isolate of SARS-CoV-2, providing a useful high-throughput BSL2 assay for studying antibody neutralization or inhibition of viral spike-mediated entry.

## RESULTS

### A replication-competent, infectious VSV chimera with SARS-CoV-2 S protein

To generate a replication-competent virus to study entry and neutralization of SARS-CoV-2 at BSL2, we engineered an infectious molecular clone of VSV by replacing the endogenous glycoprotein (G) with SARS-CoV-2 S (**Fig 1A**). SARS-CoV-2 S protein contains an endoplasmic reticulum (ER) retention sequence in the cytoplasmic tail (KxHxx-COOH) because virion assembly occurs in ER-Golgi intermediate compartments (Lontok et al., 2004; McBride et al., 2007; Ruch and Machamer, 2012). We preemptively altered that sequence to AxAxx to facilitate retargeting of S to the plasma membrane, the site of VSV assembly. Using established approaches (**Fig S1A**) (Whelan et al., 1995), we recovered infectious VSV-eGFP-SARS-CoV-2-SAA as determined by expression of the virus-encoded eGFP reporter (**Fig 1A**, *right panel*). VSV-eGFP-SARS-CoV-2-SAA propagation was inefficient on Vero CCL81 cells. This result prompted us to test additional modifications of the cytoplasmic tail of S, which were also defective in autonomous amplification (**Fig S1B**). To overcome this limitation, we used a forward genetic approach to isolate an adaptive variant of VSV-eGFP-SARS-CoV-2-SAA **(Fig S1C**). Repeated passage and plaque isolation on Vero CCL81 cells led to the emergence of a virus that contained a cysteine to stop mutation at residue 1253 (TGC to TGA at nucleotide 3759), which truncates the cytoplasmic tail of SARS-CoV-2 S by 21 residues (**Fig 1A**). We confirmed that this was the only mutation in the viral genome by next generation sequencing (Supplemental Data). Comparison of plaque morphology of VSV-eGFP-SARS-CoV-2-S_Δ21_ and VSV-eGFP-SARS-CoV-2-SAA on three Vero cell subtypes and an additional rhesus monkey cell line (MA104) demonstrates that the selected variant spreads more efficiently (**Fig 1B**). Screening of a larger panel of cell types (**Fig 1C**) identified MA104 and Vero E6 cells as supporting the highest levels of virus production. Ectopic expression of TMPRSS2 led to a further ~10-fold increase in viral titer and a larger plaque size (**Fig 1D**).

**Figure 1.**
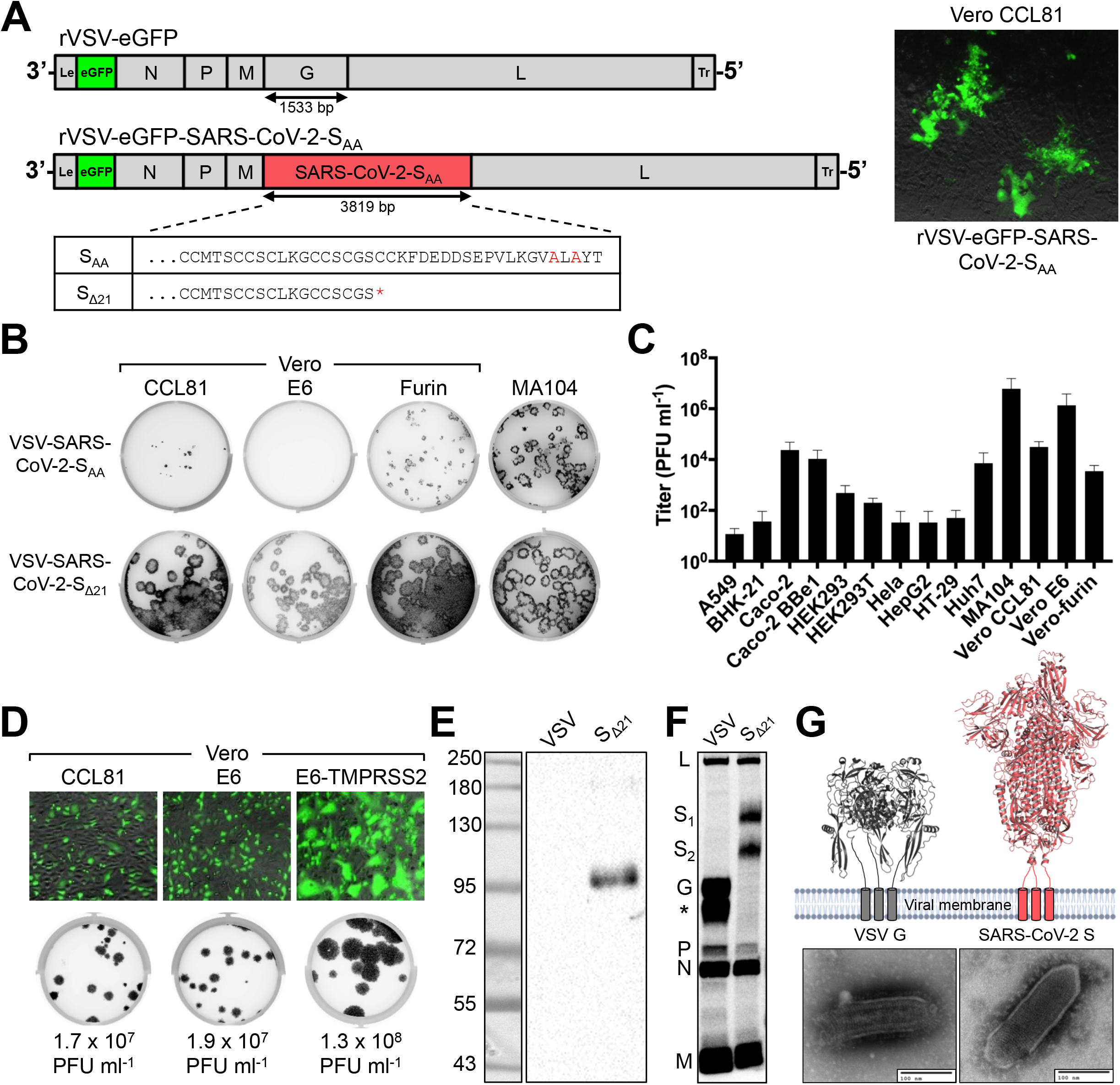
Generation and characterization of an infectious VSV-SARS-CoV-2 chimera. (**A**) A schematic diagram depicting the genomic organization of the VSV recombinants, shown 3’ to 5’ are the leader region (Le), eGFP, nucleocapsid (N), phosphoprotein (P), matrix (M), glycoprotein (G) or SARS-CoV-2 S, large polymerase (L), and trailer region (Tr). (*Right panel*) Infection of Vero CCL81 cells with supernatant from cells transfected with the eGFP reporter VSV-SARS-CoV-2-S_AA_. Images were acquired 44 hours post-infection (hpi) using a fluorescence microscope, and GFP and transmitted light images were merged using ImageJ. (*Bottom panel*) Alignment of the cytoplasmic tail of the VSV-SARS-CoV-2-S_AA_ and the sequence resulting from forward genetic selection of a mutant, which truncated the cytoplasmic tail by 21 amino acids. Mutations deviating from the wild-type spike are indicated in red, and an asterisk signifies a mutation to a stop codon. (**B**) Plaque assays were performed to compare the spread of VSV-SARS-CoV-2-S_AA_ rescue supernatant and VSV-SARS-CoV-2-S_Δ21_ on Vero CCL81, Vero E6, Vero-furin, and MA104 cells. Plates were scanned on a biomolecular imager and expression of eGFP is shown 92 hpi (representative images are shown; n>3 except for S_AA_ on Vero E6, Vero-furin, and MA104 cells). (**C**) The indicated cell types were infected with VSV-SARS-CoV-2-S_Δ21_ at an MOI of 0.5. Cells and supernatants were harvested at 24 hpi and titrated on MA104 cells (data are pooled from three or more independent experiments). (**D**) (*Top panel*) The indicated cells were infected with VSV-SARS-CoV-2-S_Δ21_ at an MOI of 2. Images were acquired 7.5 hpi using a fluorescence microscope and GFP, and transmitted light images were processed and merged using ImageJ (data are representative of two independent experiments). (*Bottom panel*) Plaque assays were performed on the indicated cell types using VSV-SARS-CoV-2-S_Δ21_. Images showing GFP expression were acquired 48 hpi using a biomolecular imager (data are representative of at least 3 independent experiments). (**E**) Western blotting was performed on concentrated VSV-SARS-CoV-2-S_Δ21_ and wild-type VSV particles on an 8% non-reducing SDS-PAGE gel. S1 was detected using a cross-reactive anti-SARS-CoV mAb (CR3022) (data are representative of two independent experiments). (**F**) BSRT7/5 cells were inoculated at an MOI of 10 with VSV-eGFP, G-complemented VSV-SARS-CoV-2-S_Δ21_, or mock infected (not shown), and metabolically labeled with [^35^S] methionine and cysteine for 20 h starting at 5 hpi in the presence of actinomycin D. Viral supernatants were analyzed by SDS-PAGE. A representative phosphor-image is shown from two independent experiments. An asterisk indicates a band that also was detected in the mock lane (not shown). (**G**) Purified VSV-WT and VSV-SARS-CoV-2-S_Δ21_ particles were subjected to negative stain electron microscopy. Prefusion structures of each respective glycoprotein are modeled above each EM image (PDB: 5I2S and 6VSB).

### SARS-CoV-2-S_Δ21_ is incorporated into infectious VSV particles

To confirm incorporation of SARS-CoV-2 S into particles, we first amplified the virus in the presence of VSV G to allow infection of cell types independently of the S protein. The VSV G *trans*-complemented VSV-SARS-CoV-2-S_Δ21_ efficiently infects HEK293T cells, which then serve as a source of production of virus particles containing SARS-CoV-2 S protein. Western blotting of supernatants with CR3022, a cross-reactive anti-S monoclonal antibody (mAb) (ter Meulen et al., 2006; Yuan et al., 2020), established the presence of S_Δ21_ in VSV-SARS-CoV-2-S_Δ21_ particles but not in the parental VSV (**Fig 1E**). The protein detected migrated at ~100 kilodaltons, a band that corresponds to the cleaved S1 subunit of the glycoprotein (Watanabe et al., 2020). To examine whether the S_Δ21_ incorporated into VSV particles is processed to S1 and S2, we performed [^35^S] cysteine-methionine metabolic labeling in BSRT7 cells, which support robust VSV replication, and analyzed released particles by SDS-PAGE and phosphorimaging. In addition to the VSV structural proteins (N, P, M and L), two additional bands were observed for VSV-SARS-CoV-2-S_Δ21_ that correspond in size to glycosylated S1 (107 kDa) and S2_Δ21_ (85 kDa) (**Fig 1F**). Negative-stain electron microscopy of sucrose-gradient purified virus particles revealed that the membrane protein projecting from VSV-SARS-CoV-2-S_Δ21_ is larger than observed on wild-type VSV particles (**Fig 1G**), which reflects the larger size of the coronavirus spike.

### A high-throughput focus-forming assay with a clinical isolate of SARS-CoV-2

VSV-SARS-CoV-2-S_Δ21_ has several advantages for detection and measuring of neutralizing antibodies, including lower biosafety containment level, ease of production and use, and rapid reporter gene readout. Nonetheless, the difference in virus morphology (spherical CoV versus bullet-shaped VSV) and possible effects on the conformational display of S on the virion surface, raise questions as to whether the accessibility of epitopes and stoichiometry of antibody neutralization is similar to authentic SARS-CoV-2. A direct comparison with a clinical isolate of SARS-CoV-2 is necessary to establish the utility of VSV-SARS-CoV-2-S_Δ21_ for assays of viral entry and antibody neutralization.

We designed a high-throughput assay for titrating SARS-CoV-2 that could be applied to multiple cell substrates. Instead of using a plaque assay, which relies on the capacity for a virus to cause cell death, which can vary across cell types, we developed a focus-forming assay (FFA) and viral antigen detection as a measure of infectivity. We propagated SARS-CoV-2 in four different producer cell types (Vero CCL81, Vero E6, Vero-furin, and MA104 cells) and then measured the number and size of foci after staining recipient cells with an anti-S mAb. With SARS-CoV-2 stocks generated from each producer cell type, we observed distinct foci across recipient cell substrates at approximately 30 h post-inoculation (**Fig 2A**). We consistently observed the highest viral titers and largest foci sizes with Vero-furin and MA104 cells (**Fig 2B-C**). However, the larger foci were more difficult to enumerate on an automated Immunospot reader and required additional manual quality control analysis. Because of this, we used Vero E6 cells for our rapid focus-reduction neutralization tests (FRNT) in subsequent experiments.

**Figure 2.**
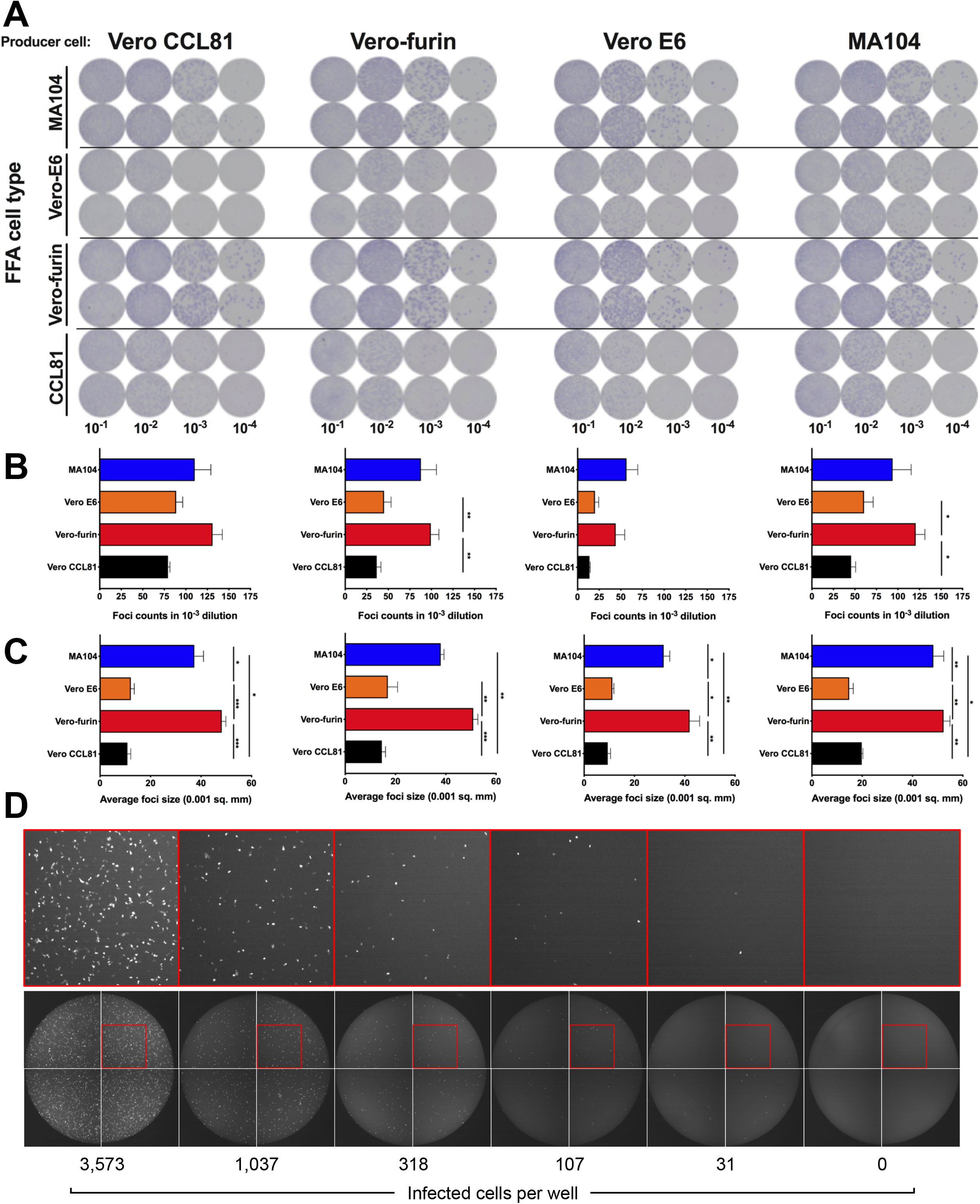
Development of a SARS-CoV-2 focus-forming assay and a VSV-SARS-CoV-2-S_Δ21_ eGFP-reduction assay. (**A**) Representative focus forming assay images of viral stocks generated from each producer cell type (top) were developed on the indicated cell substrates (indicated on the left side). Data are representative of two independent experiments. Foci obtained in (**A**) were counted (**B**) and the size was determined (**C**) using an ImmunoSpot plate reader (* *P* < 0.05, ** *P* < 0.01, *** *P* < 0.001 1 by One-way ANOVA with Tukey’s multiple comparisons test). (**D**) Representative serial dilution series of VSV-SARS-CoV-2-S_Δ21_ on Vero E6 cells. The total number of infected cells per well was quantified using an automated microscope. Insets of enhanced magnification are shown in red. Data are representative of two independent experiments.

### A high-throughput, eGFP-based neutralization assay for VSV-SARS-CoV-2-S_Δ21_

In parallel, we developed a high-throughput method to measure neutralization of VSV-SARS-CoV-2-S_Δ21_. As VSV-SARS-CoV-2-S_Δ21_ encodes an eGFP reporter and viral gene expression is robust, eGFP-positive cells can be quantified 7.5 h post-infection using a fluorescence microscope with automated counting analysis software. This approach enabled the development of an eGFP-reduction neutralization test (GRNT) (**Fig 2D**).

### Neutralization of VSV-SARS-CoV-2-S_Δ21_ and SARS-CoV-2 by human antibodies

Members of our group recently identified human mAbs from memory B cells of a SARS-CoV survivor that bind to SARS-CoV-2 S (Pinto et al, 2020). We tested a subset of these (mAbs 304, 306, 309, and 315) for their ability to inhibit VSV-SARS-CoV-2-S_Δ21_ and SARS-CoV-2 infections on Vero E6 cells. While three of these mAbs showed poor inhibitory activity, mAb 309 potently neutralized both SARS-CoV-2 and VSV-SARS-CoV-2-S_Δ21_ (**Fig 3A-B**) with similar EC50 values between the two assays (81 and 67 ng/mL for SARS-CoV-2 and VSV-SARS-CoV-2-S_Δ21_, respectively). To broaden the test panel, we evaluated the activity of a panel of mAbs generated as part of a phage display library (Sachev Sidhu, unpublished data) by both FRNT and GRNT. Many of these mAbs exhibited moderate neutralization activities in the EC50 range of 100 to 500 ng/mL (**Fig 3C-D**). Nonetheless, we observed the same neutralization trend between VSV-SARS-CoV-2-S_Δ21_ and SARS-CoV-2 with highly correlated EC50 values (< 2-fold differences).

**Figure 3.**
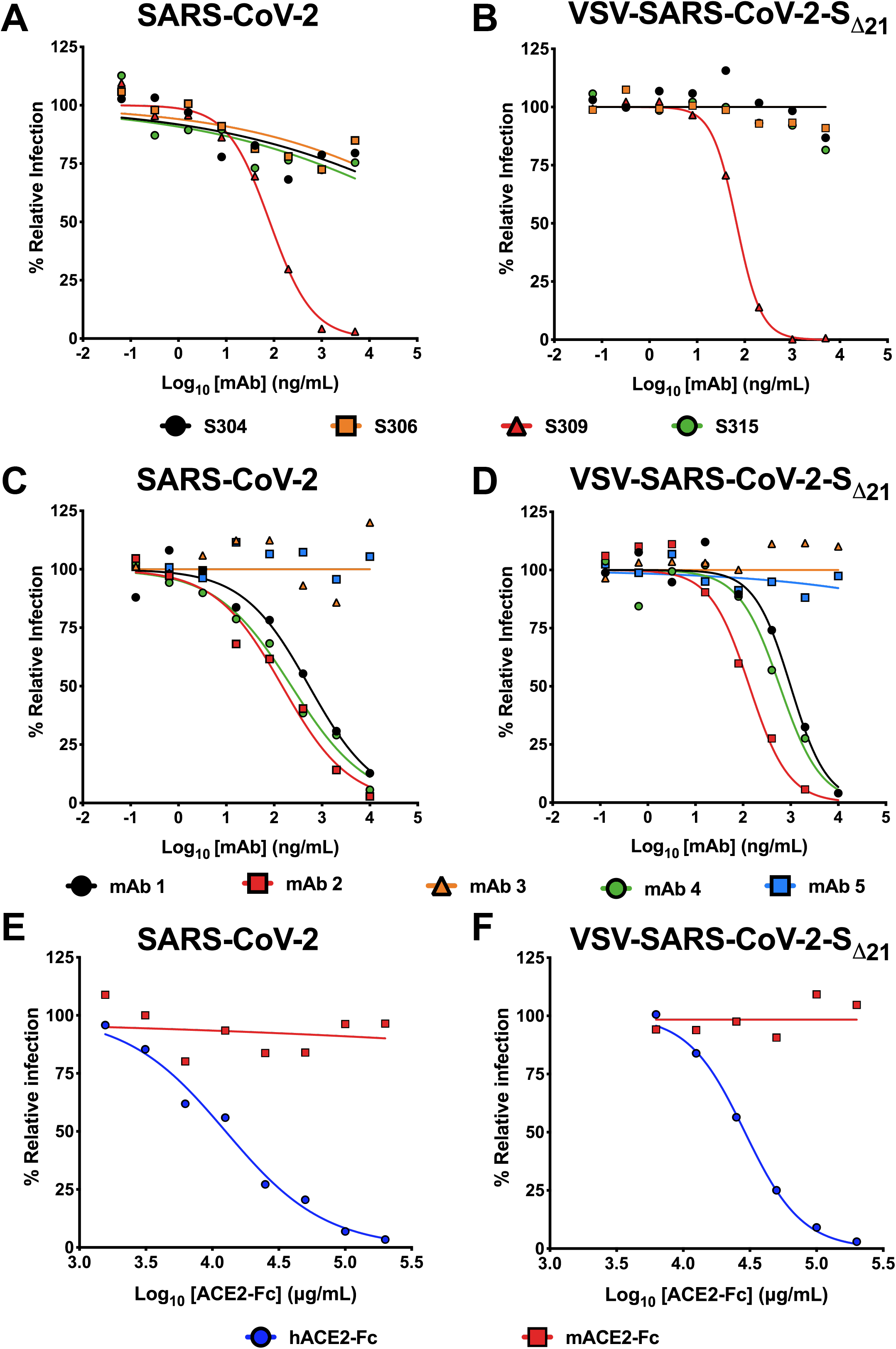
Neutralization of VSV-SARS-CoV-2-S_Δ21_ and SARS-CoV-2 by human monoclonal antibodies and hACE2 decoy receptors. **A-B.** Cross-reactive mAbs isolated from a SARS-CoV survivor were tested for neutralizing activity against SARS-CoV-2 (**A**) or VSV-SARS-CoV-2-S_Δ21_ (**B**) (n=2 and 3, respectively). **C-D**. SARS-CoV-2 RBD-specific antibodies obtained from a phage library were tested for their capacity to neutralize SARS-CoV-2 (**C**) or VSV-SARS-CoV-2-S_Δ21_ (**D**) (n=2 and 2, respectively). **E-F**. hACE2-Fc or mACE2-Fc were tested for their neutralization activity against SARS-CoV-2 (**E**) or VSV-SARS-CoV-2-S_Δ21_ (**F**) (n=2 and 3, respectively).

### Neutralization by human ACE2-Fc receptor decoy proteins

Human ACE2 (hACE2) is an entry receptor for both SARS-CoV and SARS-CoV-2 (Letko et al., 2020; Li et al., 2005; Li et al., 2003; Wrapp et al., 2020). As a soluble hACE2-Fc decoy protein has been proposed as a therapeutic for SARS-CoV-2 (Kruse, 2020), in part based on its ability to inhibit SARS-CoV infection in cell culture (Moore et al., 2004), we tested whether hACE2-Fc could inhibit infection of VSV-SARS-CoV-2-S_Δ21_ and SARS-CoV-2 using our FRNT and GRNT assays. When pre-mixed with VSV-SARS-CoV-2-S_Δ21_ or SARS-CoV-2, hACE2-Fc, but not murine ACE2-Fc (mACE2-Fc), dose-dependently and equivalently inhibited infection of recipient Vero E6 cells (**Fig 3E-F**). As expected, hACE2-Fc did not inhibit infection of wild-type VSV confirming that neutralization was specific to the SARS-CoV-2 S protein (**Fig S2**). We noted a relatively high concentration of hACE2-Fc was required for inhibition with EC50 values of 29 and 12.6 μg/ml for VSV-SARS-CoV-2-S_Δ21_ and SARS-CoV-2, respectively. Thus, although soluble hACE2-Fc decoy proteins similarly inhibit infection of VSV-SARS-CoV-2-S_Δ21_ and SARS-CoV-2, the potency is less than anticipated, which suggests that the receptor-binding domain (RBD) on the S protein on the surface of both viruses may not be fully accessible in solution.

### Neutralization of VSV-SARS-CoV-2-S_Δ21_ and SARS-CoV-2 by human immune serum

As part of studies to evaluate immune convalescent plasma as a possible therapy for SARS-CoV-2 infected patients (Bloch et al., 2020), we obtained 42 serum samples from 20 individuals at different time points after the onset of COVID-19 symptoms (**Table 1**). These samples were prescreened using a commercially available IgG ELISA. We tested each sample for neutralization of VSV-SARS-CoV-2-S_Δ21_ and SARS-CoV-2 on Vero E6 cells. We observed that sera with ELISA negative or indeterminate results generally showed low inhibitory titers (EC_50_ <1/100), whereas ELISA positive sera generated a broad range of neutralizing antibody activity (EC_50_ > 1/200 to >1/1,900) (**Fig 4A and C, Fig S3**). Remarkably, neutralization of VSV-SARS-CoV-2-S_Δ21_ and SARS-CoV-2 was similar across the entire panel of samples (**Fig 4B and C, Fig S3**).

**Table 1.** Human serum ELISA IgG. Serum samples from 20 individuals were collected at different time points post onset of COVID-19 symptoms, were screened using two ELISA assays (Euroimmun or Epitope). The serum numbers correspond to those of Figures 4 and S3. IgG index values were calculated by dividing the O.D. of the serum sample by a reference O.D. control, and ratios were interpreted using the following criteria as recommended by the manufacturer: Negative (-) <0.8, Indeterminate (+/-) 0.8-1.1, Positive (+) ≥ 1.1.

| Serum | Days post<br>symptom onset | Euroimmun IgG |  | Epitope IgG |  |
| --- | --- | --- | --- | --- | --- |
|  |  | Index | Reactive | Index | Reactive |
| 1 | 14 | 1.3 | + | 2.6 | + |
| 2 | 12 | 6.0 | + | 3.5 | + |
| 3 | 17 | 10.2 | + | 3.9 | + |
| 4 | 16 | 14.9 | + | 4.5 | + |
| 5 | 5 | 0.2 | - | 1.2 | + |
| 6 | 19 | 8.6 | + | 4.4 | + |
| 7 | 17 | 7.1 | + | 4.1 | + |
| 8 | 10 | 5.1 | + | 2.0 | + |
| 9 | 14 | 6.8 | + | 3.1 | + |
| 10 | 6 | 0.2 | - | 1.1 | + |
| 11 | - | 5.6 | + | 0.9 | +/- |
| 12 | - | <0.8 | - | 1.3 | + |
| 13 | 9 | 0.7 | - | 1.1 | + |
| 14 | 20 | 3.6 | + | 2.9 | + |
| 15 | 13 | 0.4 | - | 2.0 | + |
| 16 | 13 | 0.5 | - | 0.9 | - |
| 17 | 11 | 0.3 | - | 0.8 | - |
| 18 | 10 | 0.2 | - | 0.7 | - |
| 19 | 14 | 0.9 | +/- | 1.2 | + |
| 20 | 10 | 0.4 | - | 1.5 | + |
| 21 | 11 | 0.5 | - | 2.0 | + |
| 22 | 10 | 0.3 | - | 0.6 | - |
| 23 | 17 | 7.6 | + | 4.2 | + |
| 24 | 14 | 3.5 | + | 3.3 | + |
| 25 | 13 | 1.5 | + | 2.9 | + |
| 26 | 17 | 14.2 | + | 4.4 | + |
| 27 | 13 | 0.5 | - | 1.9 | + |
| 28 | 14 | 9.2 | + | 4.6 | + |
| 29 | 13 | 3.9 | + | 2.8 | + |
| 30 | 16 | 3.4 | + | 4.6 | + |
| 31 | 15 | 10.7 | + | 3.6 | + |
| 32 | 6 | 0.5 | - | 1.1 | + |
| 33 | 11 | 0.4 | - | 1.8 | + |
| 34 | 12 | 0.6 | - | 2.5 | + |
| 35 | 14 | 3.7 | + | 4.1 | + |
| 36 | 20 | 11.6 | + | 4.2 | + |
| 37 | 7 | 0.7 | - | 1.2 | + |
| 38 | 8 | 1.7 | + | 1.9 | + |
| 39 | 7 | 0.3 | - | 0.7 | - |
| 40 | 9 | 3.4 | + | 2.2 | + |
| 41 | 18 | 10.3 | + | 4.1 | + |
| 42 | 17 | 10.3 | + | 4.6 | + |

**Figure 4.**
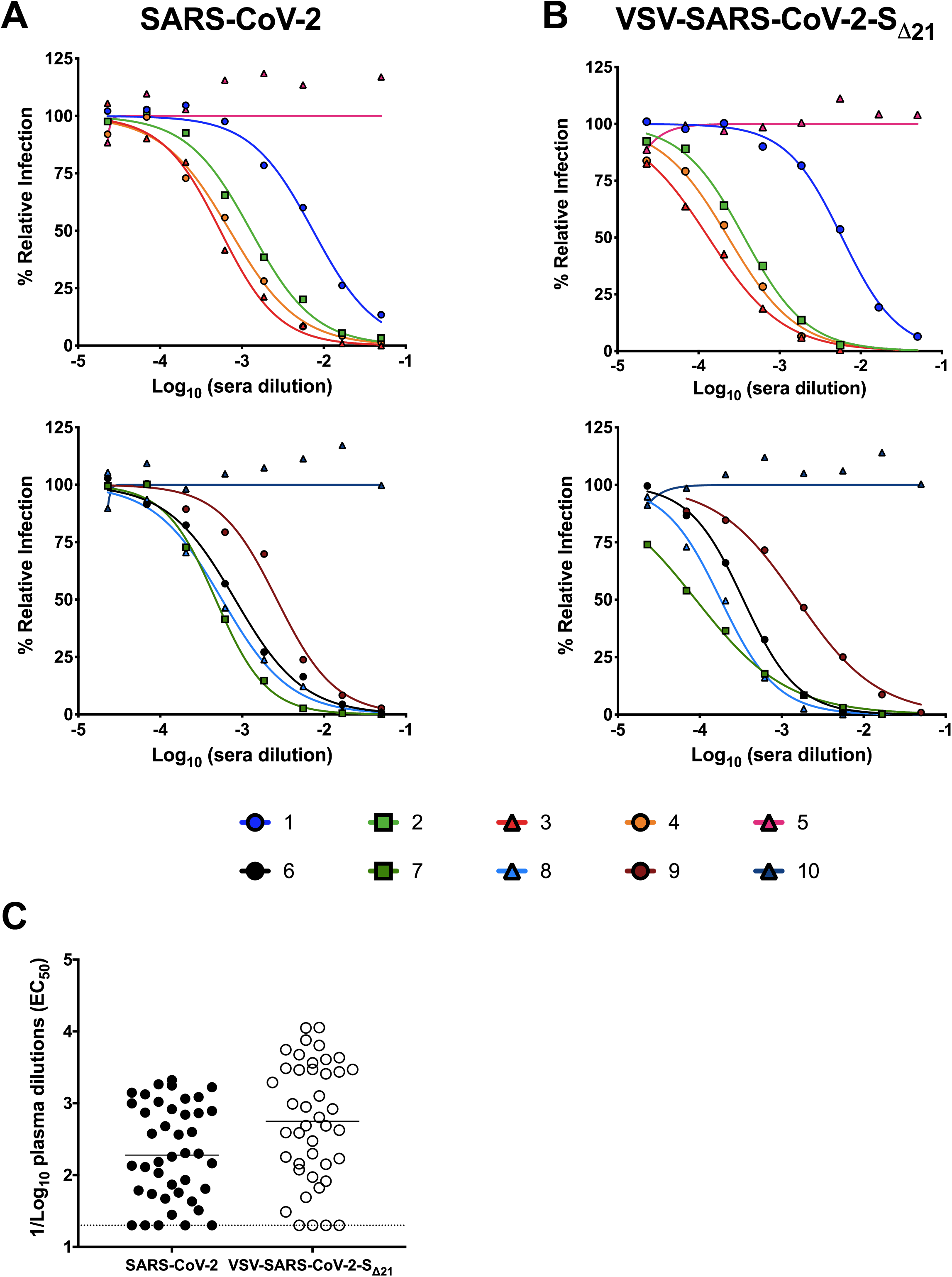
Human immune serum neutralization of SARS-CoV-2 and VSV-SARS-CoV-2-S_Δ21_. Representative neutralization curves of serum from SARS-CoV-2-infected donors with low, medium, and high inhibitory activity against SARS-CoV-2 (**A**) or VSV-SARS-CoV-2-S_Δ21_ (**B**) (n=2 and 2, respectively). (**C**) EC_50_ values of all human serum tested for neutralization of SARS-CoV-2 and VSV-SARS-CoV-2-S_Δ21_. Differences in the geometric mean or median titers were less than 3-fold between FRNT and GRNT assays.

### VSV-SARS-CoV-2-S_Δ21_ and SARS-CoV-2 neutralization assays are highly correlative

We determined the extent to which the VSV-SARS-CoV-2-S_Δ21_ and SARS-CoV-2 neutralization tests correlated with each other. We compared the GRNT and FRNT EC_50_ values obtained in assays with mAbs, polyclonal sera, and soluble ACE2 protein (**Fig 5**). For the samples with neutralizing activity, we observed a remarkably strong correlation between the two assays (*r* = 0.9285; *p* < 0.001). Moreover, all 11 of the samples that were deemed non-neutralizing in one assay had the same result in the second assay. Together, these results establish the utility of using VSV-SARS-CoV-2-S_Δ21_ as a surrogate for authentic SARS-CoV-2 infection in entry inhibition and neutralization studies.

**Figure 5.**
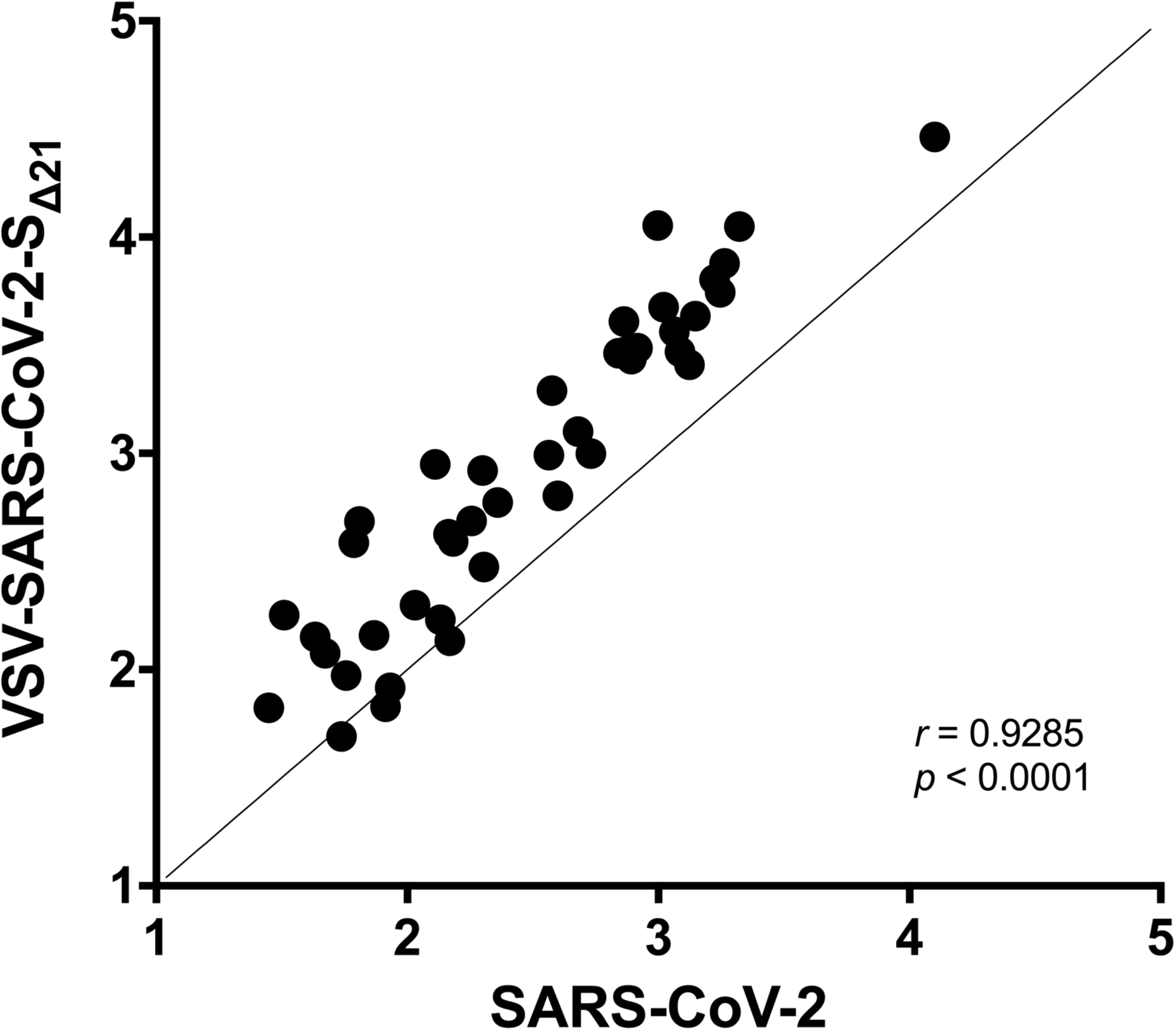
Correlation analysis of neutralization of SARS-CoV-2 and VSV-SARS-CoV-2-S_Δ21_. EC_50_ values determined in **Fig 3A-D**, and **4A-B** were used to determine correlation between neutralization assays. Spearman’s correlation *r* and *p* values are indicated.

## DISCUSSION

Emerging viral pathogens have caused numerous epidemics and several pandemics over the last century. The most recent example, SARS-CoV-2, has spread to nearly every country in the world in just a few months, causing millions of infections and hundreds of thousands of deaths (https://www.worldometers.info/coronavirus/). Rapid responses to viral outbreaks and generation of countermeasures require readily accessible tools to facilitate study and evaluate antiviral activity. Here, we generated a high-titer, replication-competent chimeric VSV expressing the SARS-CoV-2 S protein that performs similarly to a SARS-CoV-2 clinical isolate across multiple neutralization tests. As access to BSL3 facilities is limited, the finding that VSV-SARS-CoV-2-S_Δ21_ is neutralized similarly by decoy receptors, mAbs, and polyclonal antibodies in comparison to authentic SARS-CoV-2 is important. This tool will enable academic, government, and industry investigators to rapidly perform assays that interrogate SARS-CoV-2 entry, neutralization, and inhibition at a BSL2 level, which should simplify and expedite the discovery of therapeutic interventions and analysis of functional humoral immune responses.

Upon recovery of VSV-SARS-CoV-2-SAA, we selected for a mutant, which contained a 21-amino acid deletion in the cytoplasmic tail. As truncation of the cytoplasmic tail eliminates the modified KxHxx ER retention signal, we suggest that this mutation facilitates more efficient incorporation of the SARS-CoV-2 S protein into the VSV particles. Although truncation of the cytoplasmic tail of HIV envelope protein resulted in conformational alterations in the ectodomain of the protein (Chen et al., 2015), based on the extensive neutralization data presented here including correlation to neutralization of a clinical isolate of SARS CoV-2, a 21-amino acid truncation does not appear to substantively alter the structure of the S protein ectodomain. It remains to be determined whether fully wild-type S protein can incorporate efficiently into VSV. Indeed, similar mutations were generated in the SARS-CoV S protein cytoplasmic tail to boost incorporation into retroviruses and VSV pseudotypes (Fukushi et al., 2005; Giroglou et al., 2004; Moore et al., 2004).

The value of a chimeric virus depends on its capacity to present viral surface antigens in a similar way to its authentic counterpart (Garbutt et al., 2004). Indeed, the morphology of the bullet-shaped rhabdovirus and the spherical coronavirus and the density and geometry of S protein display could differentially impact antibody engagement and neutralization. Despite this concern, our extensive testing of VSV-SARS-CoV-2-S_Δ21_ with antibodies and soluble ACE2-Fc proteins showed similar neutralization profiles compared to authentic, fully infectious SARS-CoV-2. Thus, VSV-SARS-CoV-2-S_Δ21_, despite the structural differences of the virion, provides a useful tool for screening antibodies, entry-based antiviral agents, and vaccine responses against SARS-CoV-2. Indeed, convalescent plasma is under investigation as a potential COVID-19 therapeutic (Chen et al., 2020). Our studies suggest that in addition to testing for anti-S or anti-RBD antibodies (Shen et al., 2020), neutralization assays with VSV-SARS-CoV-2-S_Δ21_ may be a convenient and rapid method to obtain functional information about immune plasma preparations to enable prioritization prior to passive transfer to COVID-19 patients.

Coronaviruses possess a roughly 30 kb RNA genome, which requires that they encode a proofreading enzyme (ExoN in nsp14) (Denison et al., 2011) to counteract the error rate of the viral RNA-dependent RNA polymerase. The lack of such proofreading enzymes in the genomes of rhabdoviruses suggests that selection of escape mutants to inhibitors of the coronavirus S protein will be faster in VSV-SARS-CoV-2, which further increases the utility of this chimeric virus. Our FRNT and GRNT assays can be used to establish evidence of prior SARS-CoV-2 infection or vaccination, as well as determine waning of functional responses over time, as the likelihood of cross-neutralizing responses with other cosmopolitan coronaviruses (*e.g*., HCoV-229E and HCoV-OC43) is exceedingly low. Overall, VSV-SARS-CoV-2-S_Δ21_ and our FRNT and GRNT assays, can facilitate the development and evaluation of antibody- or entry-based countermeasures against SARS-CoV-2 infection.

## ACKNOWLEDGEMENTS

This study was supported by NIH contracts and grants (75N93019C00062, HHSN272201700060C and R01 AI127828, R37 AI059371 and U01 AI151810) and the Defense Advanced Research Project Agency (HR001117S0019) and gifts from Washington University in Saint Louis. J.B.C. is supported by a Helen Hay Whitney Foundation postdoctoral fellowship. We thank James Rini for providing RBD used to detect phage display mAbs. Some of the figures were created using BioRender.com.

## AUTHOR CONTRIBUTIONS

J.B.C. performed SARS-CoV-2 neutralization experiments. P.W.R. generated and characterized VSV-SARS-CoV-2-S_Δ21_ and performed neutralization experiments. R.E.C., Z.L., L.M.B., M.A.T., S.D., and Q.Z. provided experimental assistance. S.T., L.D., and D.W. prepared RNAseq libraries and assembled the VSV-SARS-CoV-2-S_Δ21_ sequence. H.Z. and D.H.F. generated and provided purified proteins, D.C. and H.W.V. provided the recombinant mAbs, S.M. M.U. S.S., and G.A. provided other recombinant mAbs, E.S.T. and J.P.H. identified and provided the human immune serum. M.X.G.I. helped with automated microscope use and analysis. J.B.C, P.W.R., S.P.J.W., and M.S.D. wrote the initial draft, with the other authors providing editing comments.

## SUPPLEMENTARY FIGURE LEGENDS

**Figure S1.**
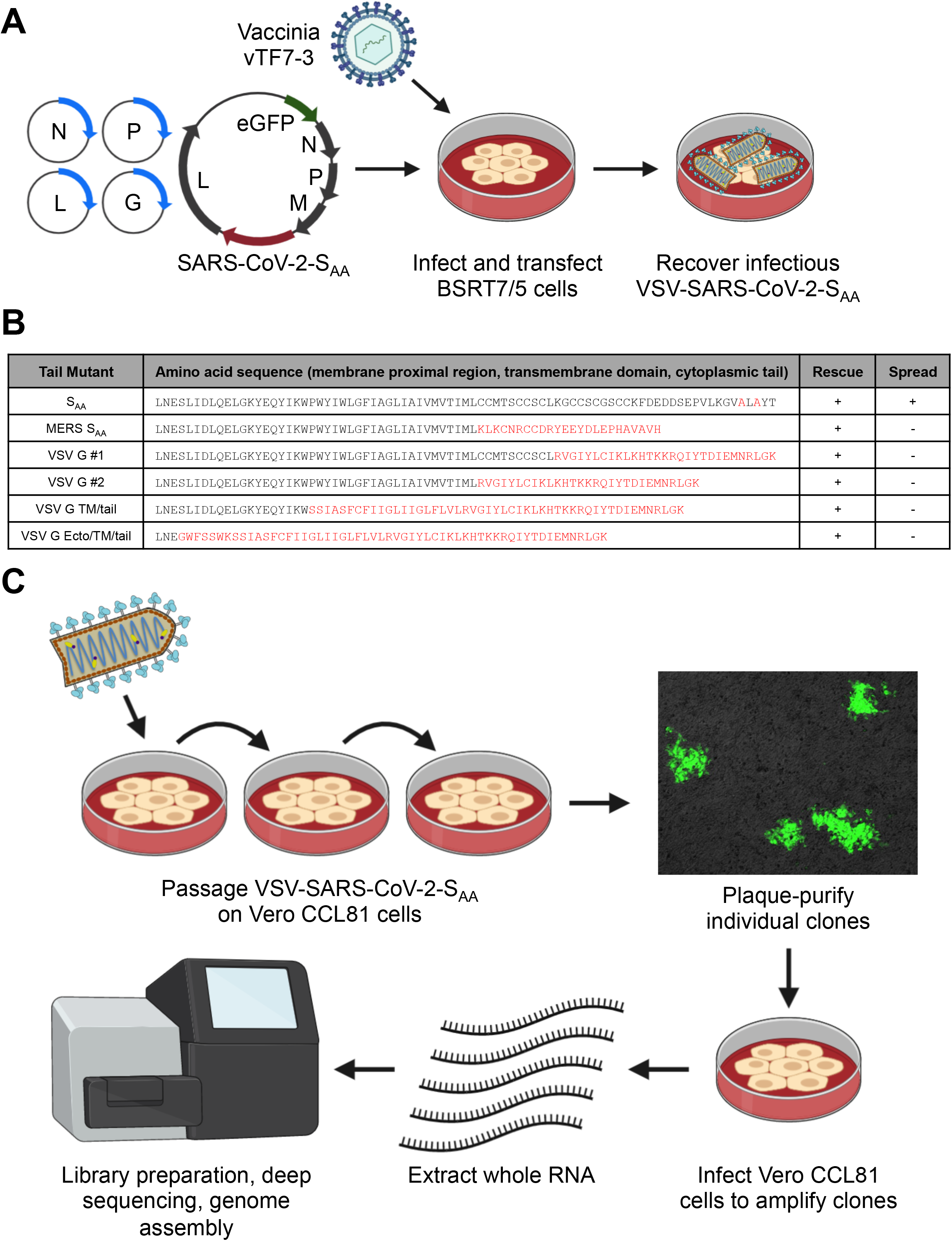
Rescue of a chimeric VSV expressing the SARS-CoV-2 S protein and forward genetic selection of a gain-of-function mutant. (**A**) BSRT7/5 cells were infected with vaccinia virus vTF7-3, transfected with plasmids allowing T7-driven expression of VSV N, P, L, and G, and an infectious molecular cDNA of VSV-SARS-CoV-2-S_AA_ to produce replication-competent VSV-SARS-CoV-2-S_AA_. (**B**) Alignment of the membrane proximal region, transmembrane domain, and cytoplasmic tail of various recombinants that were generated. Successful rescue and indication of spread are noted. (**C**) VSV-SARS-CoV-2-SAA was passaged iteratively on Vero CCL81 cells. Several clones were plaque-purified and amplified on Vero CCL81 cells. RNA from infected cells was extracted and deep sequenced to identify mutants.

**Figure S2.**
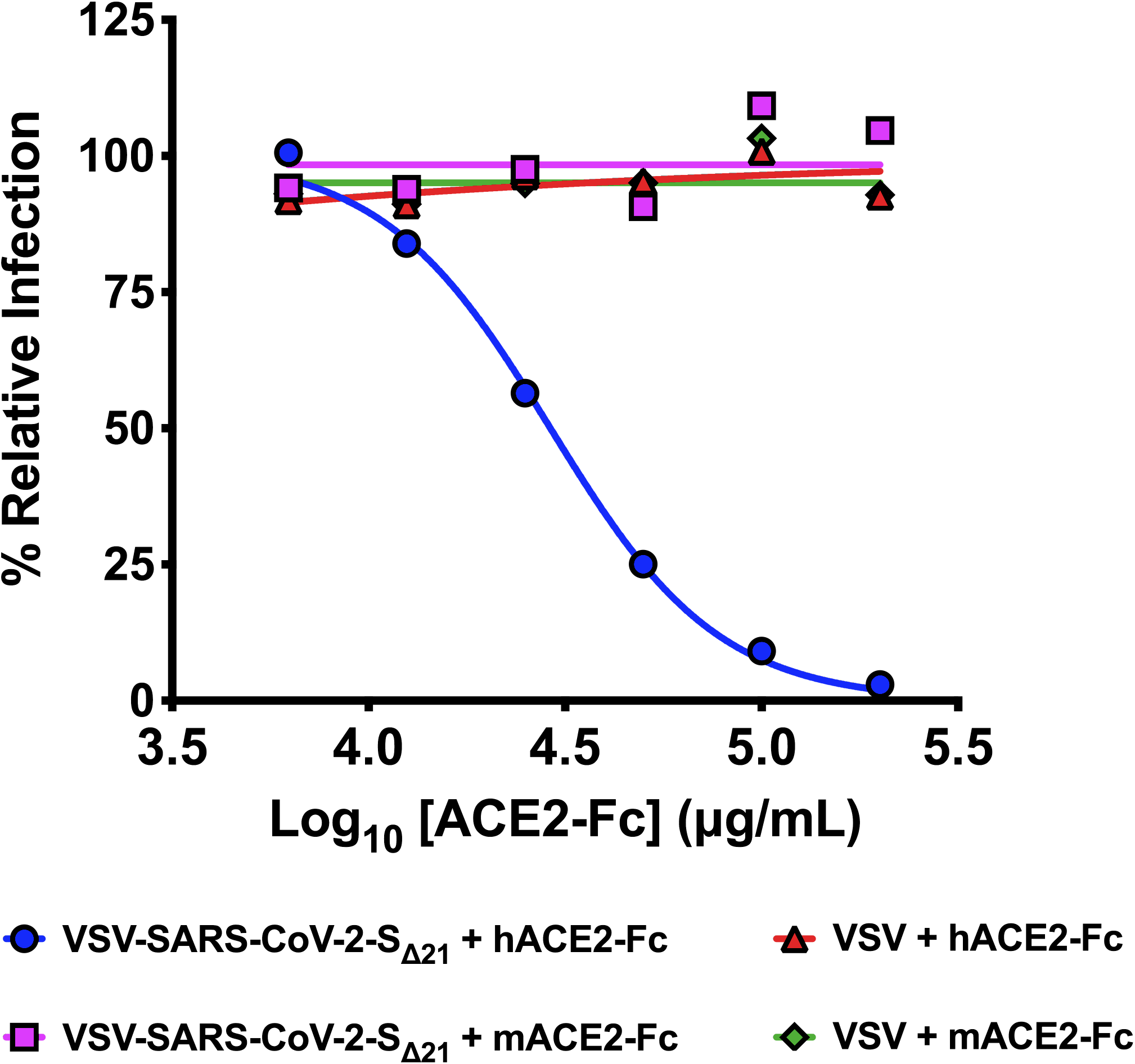
Inhibition of VSV-SARS-CoV-2-S_Δ21_ but not VSV with hACE2-Fc receptor decoy proteins. VSV-SARS-CoV-2-S_Δ21_ and VSV were incubated with the indicated human or murine ACE2-Fc receptor decoy proteins, and virus-antibody mixtures were used to infect Vero E6 cells in a GRNT assay. Data are representative of three independent experiments.

**Figure S3.**
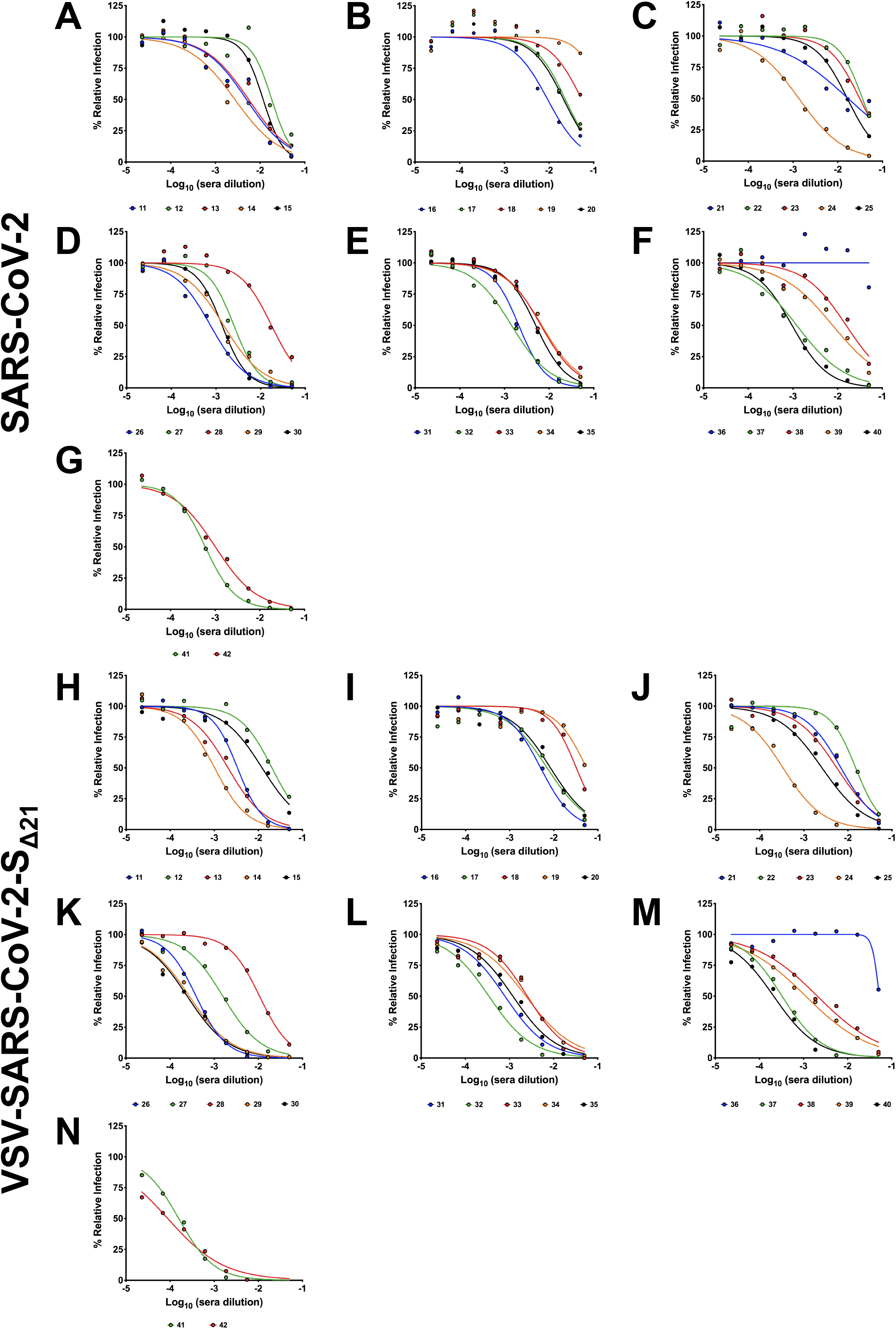
Human immune serum neutralization of SARS-CoV-2 and VSV-SARS-CoV-2-S_Δ21_. As described in **Fig 4**, human serum samples from PCR confirmed SARS-CoV-2-infected patients were tested in FRNT (**A-G**) and GRNT (**H-N**) assays with SARS-CoV-2 and VSV-SARS-CoV-2-S_Δ21_.

## METHODS

### Plasmids

The S gene of SARS-CoV-2 isolate Wuhan-Hu-1 (GenBank MN908947.3) was synthesized in two fragments (Integrated DNA Technologies) and inserted into an infectious molecular clone of VSV (Whelan et al., 1995) as previously (Carette et al., 2011; Jae et al., 2014). Modifications to the cytoplasmic tail were assembled identically. Other plasmids were previously described: VSV N, P, L and G expression plasmids (Stanifer et al., 2011; Whelan et al., 1995), psPAX2 (Addgene), and pLX304-TMPRSS2 (Zang et al., 2020).

### Cells

Cells were maintained in humidified incubators at 34 or 37°C and 5% CO2 in the indicated media. BSRT7/5, Vero CCL81, Vero E6, Vero E6-TMPRSS2, A549, Caco-2, Caco-2 BBe1, Huh7, HepG2, Hela, BHK-21, HEK293, and HEK293T were maintained in DMEM (Corning or VWR) supplemented with glucose, L-glutamine, sodium pyruvate, and 10% fetal bovine serum (FBS). Vero-furin cells (Mukherjee et al., 2016) also were supplemented with 5 μg/ml of blasticidin. MA104 cells were maintained in Medium 199 (Gibco) containing 10% FBS. HT-29 cells were cultured in complete DMEM/F12 (Thermo-Fisher) supplemented with sodium pyruvate, non-essential amino acids, and HEPES. Vero E6-TMPRSS2 cells were generated using a lentivirus vector. Briefly, HEK293T producer cells were transfected with pLX304-TMPRSS2, pCAGGS-VSV-G, and psPAX2, and cell culture supernatants were collected at 48 hours and clarified by centrifugation at 1,000 x g for 5 min. The resulting lentivirus was used to infect Vero E6 cells for 24 h, and cells were selected with 40 μg/ml blasticidin for 7 days.

### Recombinant VSV

Recovery of recombinant VSV was performed as described (Whelan et al., 1995). Briefly, BSRT7/5 cells were inoculated with vaccinia virus vTF7-3 and subsequently transfected with T7-expression plasmids encoding VSV N, P, L, and G, and an antigenomic copy of the viral genome. Cell culture supernatants were collected at 56-72 h, clarified by centrifugation (5 min at 1,000 x g), and filtered through a 0.22 μm filter. Virus was plaque-purified on Vero CCL81 cells in the presence of 25 μg/ml of cytosine arabinoside (TriLink BioTechnologies), and plaques in agarose plugs were amplified on Vero CCL81 cells. Viral stocks were amplified on MA104 cells at an MOI of 0.01 in Medium 199 containing 2% FBS and 20 mM HEPES pH 7.7 (Millipore Sigma) at 34°C. Viral supernatants were harvested upon extensive cytopathic effect and clarified of cell debris by centrifugation at 1,000 x g for 5 min. Aliquots were maintained at −80°C.

### Next generation sequencing

Total RNA was extracted from Vero CCL81 cells infected with VSV-SARS-CoV-2-S_Δ21_ using Trizol (Invitrogen) according to the manufacturer’s protocol. RNA was used to generate next generation sequencing libraries using TruSeq Stranded Total RNA library kit with Ribo Zero ribosomal subtraction (Illumina). The libraries were quantified using a bioanalyzer (Agilent) and pooled at an equal molar concentration and used to generate paired end 250 bp reads on an MiSeq (Illumina). Raw sequencing data was processed using fastp v0.20.0 to trim adapters and filter out reads with a quality score < 30. Alignment of each sample to VSV and SARS CoV-2 sequence was performed using bbmap v38.79. Mapped reads were extracted using samtools 1.9 and used for de novo assembly by SPAdes v3.13.0. Consensus sequences for each RNA sample were generated by aligning contigs to the reference plasmid sequence pVSV(1+)-eGFP-SARS-CoV-2-S with SnapGene v5.0.

### Western blotting

Purified VSV were incubated in non-reducing denaturation buffer (55 mM Tris-HCl pH 6.8, 1.67 % (w/v) SDS, 7.5 % (w/v) glycerol) at 100°C for 5 min. Viral proteins were separated on a 8% acrylamide gel, transferred onto a nitrocellulose membrane, and incubated with human anti-SARS antibody CR3022 diluted in Tris-buffer saline containing 1% Tween-20 (TBS-T) and 5% milk, followed by incubation with HRP-conjugated goat anti-human antibody (Abcam) diluted in TBS-T containing 1% milk. HRP activity was visualized using the Pierce ECL Western blotting kit (Thermo Scientific) and imaged with a ChemiDocTM MP Imager (Bio-Rad).

### Metabolic radiolabeling of virions

To generate high titer stocks of VSV-SARS-CoV-2, BSRT7/5 cells were transfected with pCAGGS-VSV-G in Opti-MEM (Gibco) using Lipofectamine 2000 (Invitrogen) and infected 7 h later with VSV-SARS-CoV-2-S_Δ21_ at an MOI of 0.1 in DMEM containing 2% FBS and 20 mM HEPES pH 7.7. Viral stocks were collected at 48 hpi, and used to infect fresh cells (MOI of 10) for labeling of viral proteins. At 4 hpi, cells were starved in serum free, methionine/cysteine free DMEM (Corning), and exposed to 15 μCi/ml [^35^S]-methionine and [^35^S]-cysteine (Perkin Elmer) from 5-24 hours. Cell culture supernatants were collected, clarified by centrifugation (1,500 x g, 5 min), and analyzed by SDS-PAGE and detected by phosphoimage analysis (Li et al., 2006).

### Transmission electron microscopy

Purified viruses were adhered to glow-discharged, carbon-coated copper grids. Samples were stained with 2% (w/v) phosphotungstic acid, pH 7.1, in H2O and viewed on a JEOL 1200 EX transmission electron microscope (JEOL USA Inc.) equipped with an AMT 8-megapixel digital camera and AMT Image Capture Engine V602 software (Advanced Microscopy Techniques).

### Monoclonal antibodies

Phage-displayed Fab libraries were panned against immobilized SARS-CoV-2 spike RBD in multiple rounds using established methods (Persson et al., 2013). Following four rounds of selection, phage ELISAs were used to screen 384 clones to identify those that bound specifically to RBD. The complementarity determining regions of Fab-phage clones were decoded by sequencing the variable regions and cloning them into mammalian expression vectors for expression and purification of human IgG1 proteins, as described (Tao et al., 2019). A subset of the panel of mAbs was tested for neutralization as a part of this study.

Another set of mAbs (S304, S306, S309, S310 and S315) were isolated from EBV-immortalized memory B cells from a SARS-CoV survivor (Traggiai et al., 2004) and are cross-reactive to SARS-CoV-2 (Pinto et al, in press). Recombinant antibodies were expressed in ExpiCHO cells transiently co-transfected with plasmids expressing the heavy and light chain as previously described (Stettler et al., 2016).

### Human sera

Human samples were collected from PCR-confirmed COVID-19 patients. Serum samples were obtained by routine phlebotomy at different days post symptom onset (range: day 5 - 20). Samples were prescreened by the Euroimmun anti-SARS-CoV-2 IgG ELISA (Lubeck, Germany), a qualitative assay with the Food and Drug Administration Emergency Use Authorization that detects antibodies to the SARS-CoV-2 S protein. This study was approved by the Mayo Clinic Institutional Review Board.

### Protein expression and purification

DNA fragments encoding human ACE2 (hACE2 residues 1-615) and mouse ACE2 (mACE2, residues 1-615) were synthesized and cloned into pFM1.2 with a C-terminal HRV-3C protease cleavage site (LEVLFQGP) and a human IgG1 Fc region as previously described (Raj et al., 2013). We transiently transfected plasmids into Expi293F cells and harvested cell supernatants 4 days post transfection. Secreted hACE2-Fc and mACE2-Fc proteins were purified by protein A chromatography (Goldbio).

### GFP-reduction neutralization test

Patient samples were heat-inactivated at 56°C for 30 min. Indicated dilutions of samples were incubated with 10^2^ PFU of VSV-SARS-CoV-2-S_Δ21_ for 1 h at 37°C. Antibody-virus complexes were added to Vero E6 cells in 96-well plates and incubated at 37°C for 7.5 h. Cells subsequently were fixed in 2% formaldehyde containing 10 μg/mL Hoechst 33342 nuclear stain (Invitrogen) for 45 min at room temperature, when fixative was replaced with PBS. Images were acquired with the InCell 2000 Analyzer (GE Healthcare) automated microscope in both the DAPI and FITC channels to visualize nuclei and infected cells (*i.e*., eGFP-positive cells), respectively (4X objective, 4 fields per well, covering the entire well). Images were analyzed using the Multi Target Analysis Module of the InCell Analyzer 1000 Workstation Software (GE Healthcare). GFP-positive cells were identified in the FITC channel using the top-hat segmentation method and subsequently counted within the InCell Workstation software. The sensitivity and accuracy of GFP-positive cell number determinations were validated using a serial dilution of virus. A background number of GFP+ cells was subtracted from each well using an average value determined from at least 4 uninfected wells. Data were processed using Prism software (GraphPad Prism 8.0). ACE2 neutralization assays using VSV-SARS-CoV-2-S_Δ21_ were conducted similarly but using an MOI of 1.

### Focus reduction neutralization test

SARS-CoV-2 strain 2019 n-CoV/USA_WA1/2020 was obtained from the Centers for Disease Control and Prevention (gift of Natalie Thornburg). Virus was passaged in the indicated producer cells. Indicated dilutions of mAbs, sera, or protein were incubated with 10^2^ FFU of SARS-CoV-2 for 1 h at 37°C. Antibody-virus complexes were added to indicated cell monolayers in 96-well plates and incubated at 37°C for 1 h. Subsequently, cells were overlaid with 1% (w/v) methylcellulose in MEM supplemented with 2% FBS. Plates were harvested 30 h later by removing overlays and fixed with 4% paraformaldehyde in PBS for 20 min at room temperature. Plates were washed and sequentially incubated with 1 μg/mL of CR3022 anti-S antibody (ter Meulen et al., 2006; Yuan et al., 2020) and HRP-conjugated goat anti-human IgG in PBS supplemented with 0.1% saponin and 0.1% BSA. SARS-CoV-2-infected cell foci were visualized using TrueBlue peroxidase substrate (KPL) and quantitated on an ImmunoSpot microanalyzer (Cellular Technologies). Data were processed using Prism software (GraphPad Prism 8.0).

### Statistical analyses

All statistical tests were performed using GraphPad Prism 8.0 software as described in the indicated figure legends.

